# A feasibility study of smartphone sensors to assess the effect of acute high altitude (3,800 m) exposure on cognition and motor function in healthy participants

**DOI:** 10.1101/2023.10.09.561578

**Authors:** Oliver G. Goldman, Gerald Dubowitz, David Anderson

## Abstract

Acute exposure to hypoxia at attitude has neurologic effects. Some subjects develop severe neurologic symptoms, including Parkinsonism, when hypoxic at high altitude as part of an Acute Mountain Sickness syndrome. Digital health technologies can provide continuous monitoring and objective, real-world measures of movement disorders, but only limited validation data of wearable sensors is available in the high-altitude setting. This equipment validation and feasibility study assessed smartphone microphone and accelerometer function at sea level and 12470 feet (3,800 m) to assess their potential value to predict and prevent neurologic features of acute mountain sickness. A feasibility study of standardized assessments of motor, speech and cognitive tasks was performed in 3 normal subjects at sea level and at altitude. All subjects were hypoxic at altitude with O_2_ saturations ranging from 77-81%. Shaker table (range of frequencies) and high-fidelity speaker (range of frequencies) controls confirmed high correlation of observed and expected measurements for microphone and accelerometer under all conditions. The feasibility study demonstrated that under conditions of hypoxia at attitude, fine motor skills are impaired; visual short-term memory is not impaired but has longer response time; gait and balance is impaired, and a postural tremor develops with frequencies below 10 Hz. Future studies could use these wearable sensors to further assess effects at altitude of more severe hypoxia with applications in the high-altitude environment for Parkinson’s Disease patients, with further opportunity for aviation and military use.

## INTRODUCTION

Acute mountain sickness (AMS) is characterized by effects on motor function and cognition [1]. These include impairment of memory, fine motor function, balance, and gait [2]. Parkinson’s symptoms have been reported at altitude in hypoxic subjects with and without a history of movement disorder [3, 4]. Prediction and prevention of AMS is challenging due to the previous lack of sensitive equipment in the remote environment, indicating the potential benefits of digital tools. These tools provide sensitive, real-world measures of movement disorders such as Parkinson’s Disease [5];[6] [7]; [8]. A smartphone research application, previously measured tremor and detected motor fluctuations and dyskinesias with high correlation to clinical disease at sea-level, but the sensors have not been validated at altitude [9, 10].

If validated for use at altitude, sensitive consumer wearables could be used as predictive and preventive tools in the assessment of AMS and in understanding the vulnerability of Parkinson’s Disease patients to deterioration at altitude.

## METHODS

### Participants and Setting

All equipment validation and behavior feasibility experiments were conducted at sea level and at immediately on arrival at (12,470 feet; 3,800 m). Participants were all male, aged 17, 57, 58 years with no concomitant medications and no relevant past medical history. All altitude recordings were performed immediately on arrival at University of California White Mountain Research center (WMRC), a High Altitude research facility (https://www.wmrc.edu). All participants had hypoxia verified using WMRC reference oximeters (O_2_ sat 77-81% at altitude). None of the participants had acute mountain sickness as measured by the Lake Louise score [11].

### Equipment Validation Studies

We used a commercially available tabletop shaker (Serology Rotator, LW Scientific). The shaker was set to rotational speeds of 1, 2, 3, and 4 Hz (rotations per second). Accelerometer output (iPhone 14) was evaluated in 3 dimensions. Correspondence between expected and observed frequencies was assessed. A smartphone (iPhone 14) was placed near a high-fidelity speaker (JBL PartyBox 100). Standardized tones of 50, 100, 150, 200, 250, and 300 Hz were played. Speaker output was recorded using smartphone microphone. Correspondence between expected and observed frequencies was evaluated.

### Evaluating Cognition and Mobility

An Apple Watch 8 and an iPhone 14 (Apple, Inc.) running a smartphone application specifically designed by Clinical ink for movement disorders (BrainBaseline ™) was used. Raw mobility and speech signals were recorded from Apple’s native accelerometer (100 Hz sampling rate) and microphone (32 kHz sampling rate) hardware configurations. Smartphone application tasks were conducted at sea-level and altitude. The smartphone was worn in a lumbar sport pouch during gait and balance tests. Gait features were extracted from the smartwatch and smartphone using software developed in-house. Participants wore the smartwatch and accelerometry data and tremor scores were collected from the smartwatch via Apple’s Movement Disorders Application Programming Interface. Tremor analysis was performed on participants with at least 24 h of passive data over two weeks after baseline. Using the BrainBaseline™ App, a cognitive and psychomotor battery was administered via the smartphone that included the Trail Making Test, modified Symbol Digit Modalities Test, Visuospatial Working Memory Task, and two timed fine motor tests [12]; [13] [14].

Speech tasks included articulation, phonation and reading. These files were processed using custom Python code (available from the authors upon request) with features computed using the Librosa library [15]. Common speech endpoints, such as jitter, shimmer, pitch statistics were computed. Speech segmentation was performed and used to extract time-related features for reading tasks [16]

## RESULTS

### Accelerometer Validation

Fourier analysis of raw accelerometer output produced by shaker table movement at both sea level (Figure 1A) and at altitude (Figure 1B) revealed clear peaks in frequencies programmed into the shaker table during each trial. As expected, no clear activity was observed in the z-axis, corresponding to accelerometer axis perpendicular to the plane of the shaker table surface. Monotonic increases in peak frequency amplitudes as a function of shaker table rotational frequency reflect increasing velocity of the table at higher rotational frequencies. Plots of observed peak frequencies as a function of expected peak frequencies demonstrated good correspondence for x- and y-axes at both sea level (Figure 1C; r_x_=0.99994; r_y_=0.99994) and altitude (Figure 1D; r_x_=0.99994; r_y_=0.99994).

**Figure 1.**
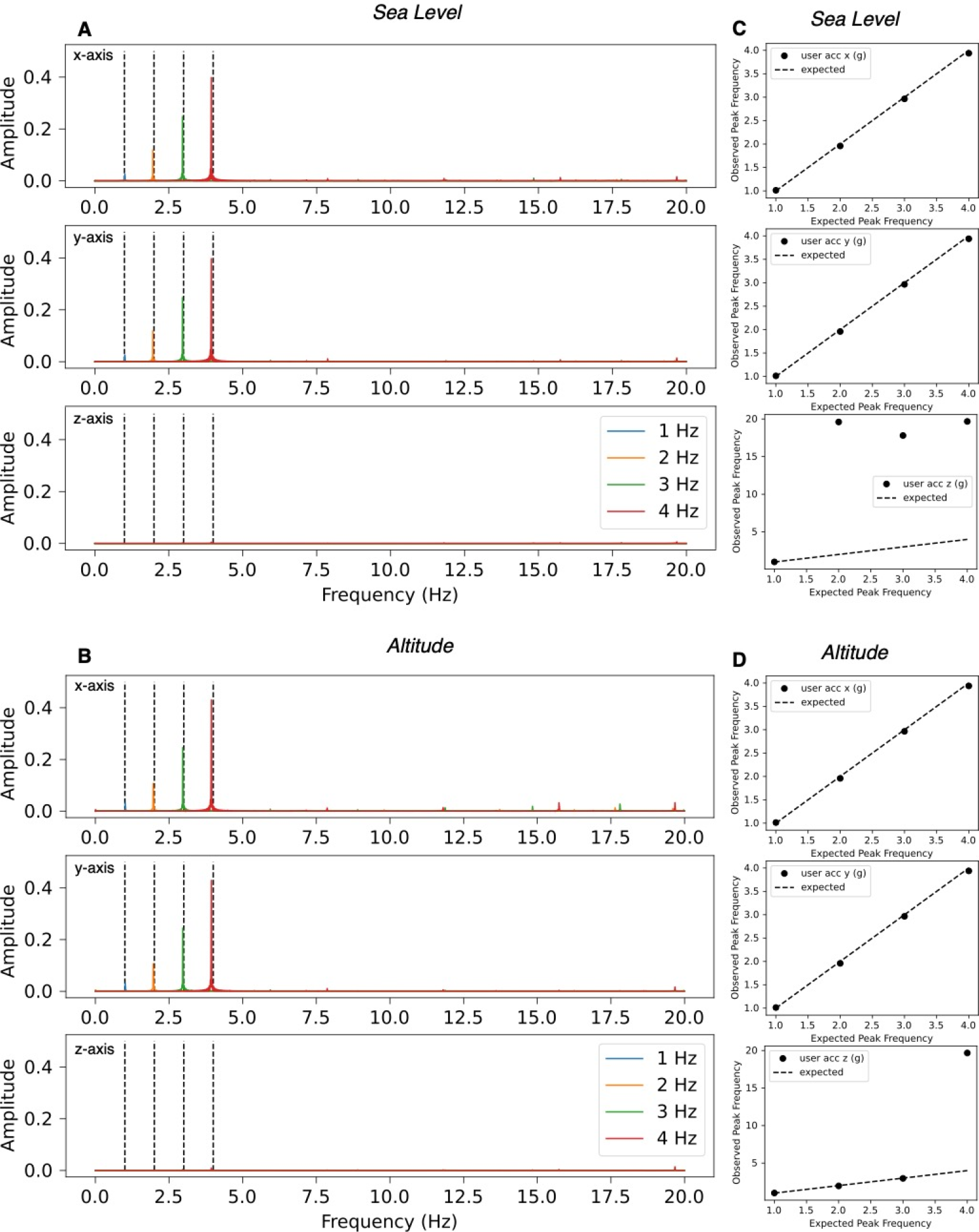
Accelerometer Validation. **(A-B)** Spectral activity derived from Fourier transform of raw accelerometer data generated from a range of rotational frequencies at sea level **(A)** and at altitude **(B)**. Each colored line corresponds to a unique frequency. Dotted lines correspond to the frequencies programmed into the shaker table. Each plot corresponds to a different axis of the tri-dimensional accelerometer. **(C-D)**. Input-output plots of expected and observed peak frequencies observed at sea level **(C)** and at altitude **(D)**. Each plot corresponds to a different axis. Dotted lines correspond to the expected frequency.

### Microphone Validation

Fourier analysis of audio waveforms produced by the microphone in response to pure tones at both sea level (Figure 2A) and at altitude (Figure 2B) revealed clear peaks in frequencies represented in the pure tones on each trial. Plots of observed peak frequencies as a function of expected peak frequencies demonstrated good correspondence at both sea level (Figure 1C; r=0.99999990) and altitude (Figure 1D; r=0.9999995).

**Figure 2.**
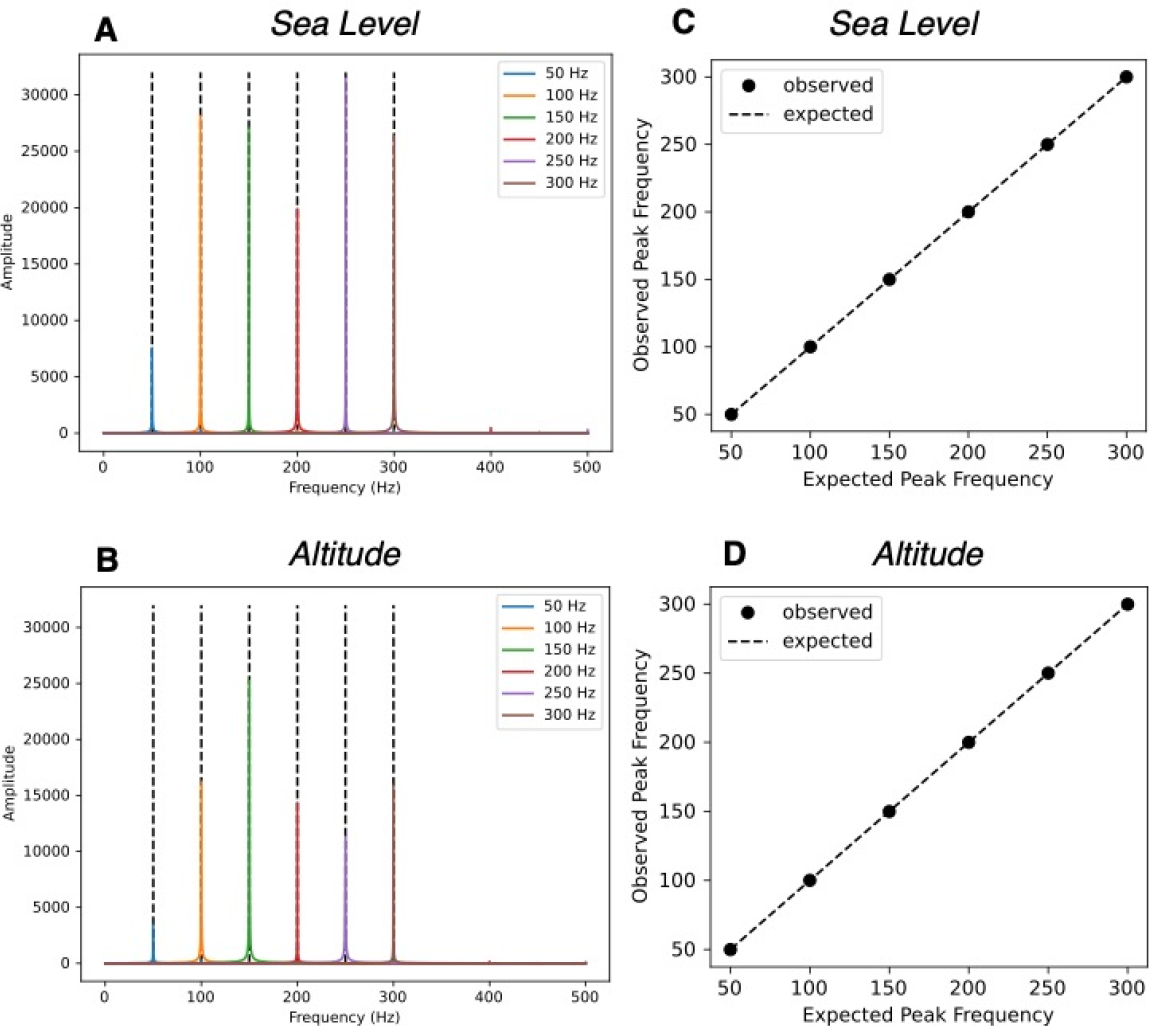
Microphone Validation. **(A-B)** Spectral activity derived from Fourier transform of microphone data generated from a range of pure tones at sea level **(A)** and at altitude **(B)**. Each colored line corresponds to a unique frequency. Dotted lines correspond to the frequencies within each tone. **(C-D)**. Input-output plots of expected and observed peak frequencies observed at sea level **(C)** and at altitude **(D)**. Dotted lines correspond to the expected frequency.

### Evaluating Cognition and Mobility

Participants completed all assessments on the application twice in both settings. Endpoints generated from each trial are summarized in Table 1. To minimize contributions of practice effects, preliminary descriptive analyses focused only on the second trial from each environment. At 3,800 m altitude the following descriptive trends were noted:

**Table 1.**
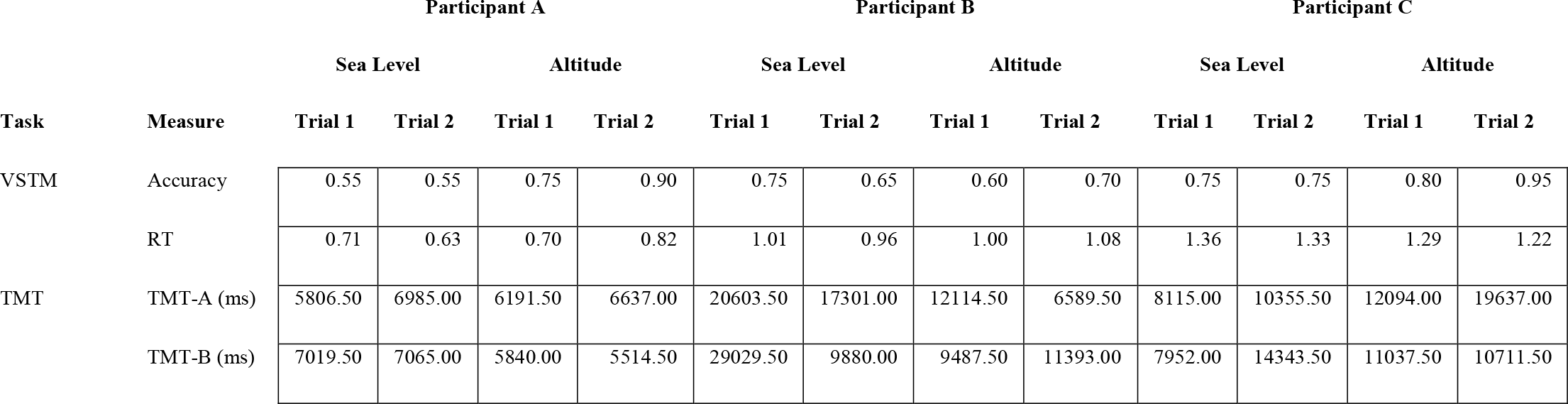
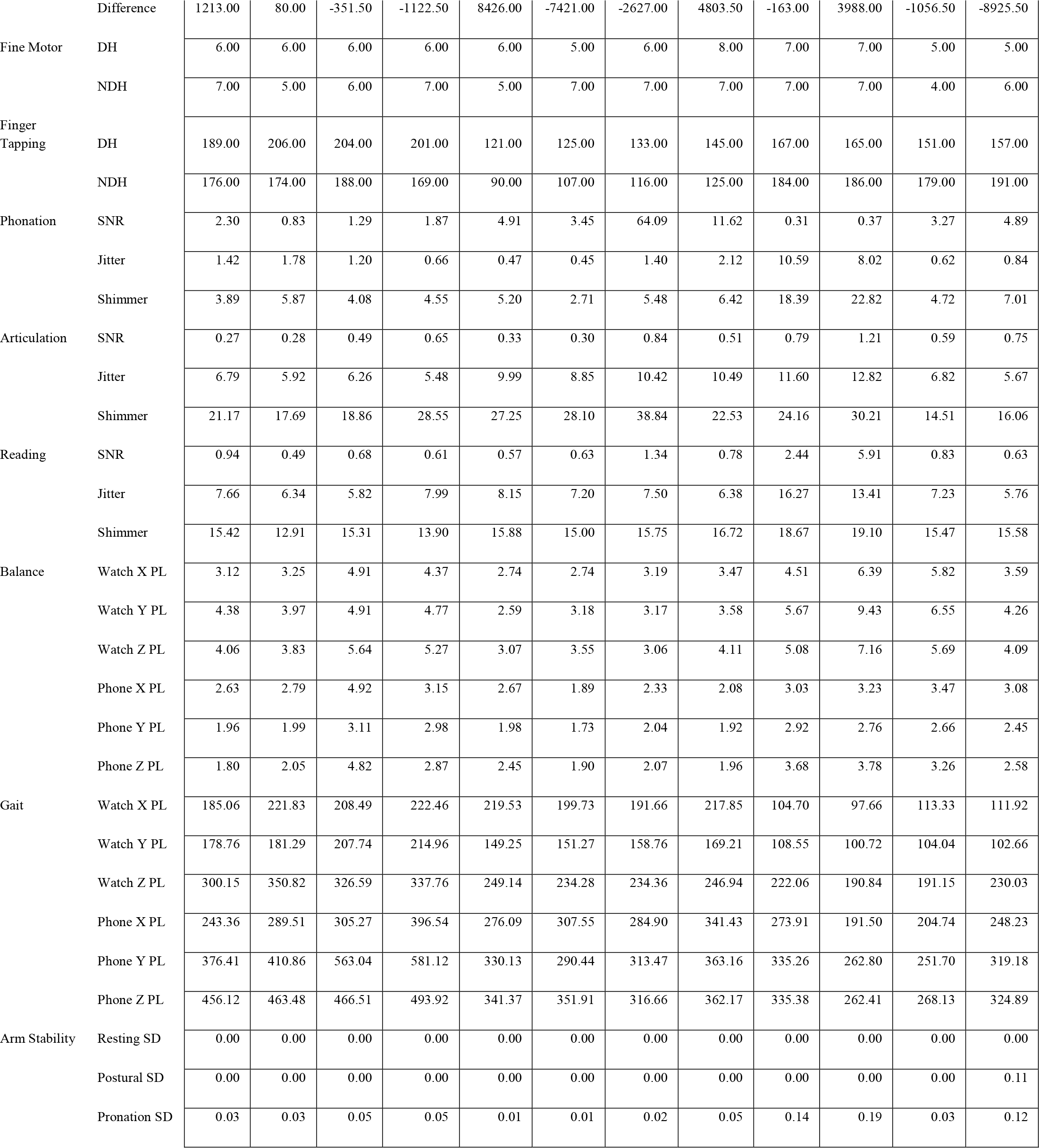
Summary of all endpoints generated by each participant at sea level and at altitude for both trials. (DH = dominant hand; NDH = non-dominant hand; PL = path length; RT = reaction time; SD = standard deviation; SNR = signal-to-noise ratio; TMT = trail making test; VSTM=visual short-term memory).

| Task | Measure | Participant A |  |  |  | Participant B |  |  |  | Participant C |  |  |  |
| --- | --- | --- | --- | --- | --- | --- | --- | --- | --- | --- | --- | --- | --- |
|  |  | Sea Level |  | Altitude |  | Sea Level |  | Altitude |  | Sea Level |  | Altitude |  |
|  |  | Trial 1 | Trial 2 | Trial 1 | Trial 2 | Trial 1 | Trial 2 | Trial 1 | Trial 2 | Trial 1 | Trial 2 | Trial 1 | Trial 2 |
| VSTM | Accuracy | 0.55 | 0.55 | 0.75 | 0.90 | 0.75 | 0.65 | 0.60 | 0.70 | 0.75 | 0.75 | 0.80 | 0.95 |
|  | RT | 0.71 | 0.63 | 0.70 | 0.82 | 1.01 | 0.96 | 1.00 | 1.08 | 1.36 | 1.33 | 1.29 | 1.22 |
| TMT | TMT-A (ms) | 5806.50 | 6985.00 | 6191.50 | 6637.00 | 20603.50 | 17301.00 | 12114.50 | 6589.50 | 8115.00 | 10355.50 | 12094.00 | 19637.00 |
|  | TMT-B (ms) | 7019.50 | 7065.00 | 5840.00 | 5514.50 | 29029.50 | 9880.00 | 9487.50 | 11393.00 | 7952.00 | 14343.50 | 11037.50 | 10711.50 |
|  | Difference | 1213.00 | 80.00 | -351.50 | -1122.50 | 8426.00 | -7421.00 | -2627.00 | 4803.50 | -163.00 | 3988.00 | -1056.50 | -8925.50 |
| Fine Motor | DH | 6.00 | 6.00 | 6.00 | 6.00 | 6.00 | 5.00 | 6.00 | 8.00 | 7.00 | 7.00 | 5.00 | 5.00 |
|  | NDH | 7.00 | 5.00 | 6.00 | 7.00 | 5.00 | 7.00 | 7.00 | 7.00 | 7.00 | 7.00 | 4.00 | 6.00 |
| Finger Tapping | DH | 189.00 | 206.00 | 204.00 | 201.00 | 121.00 | 125.00 | 133.00 | 145.00 | 167.00 | 165.00 | 151.00 | 157.00 |
|  | NDH | 176.00 | 174.00 | 188.00 | 169.00 | 90.00 | 107.00 | 116.00 | 125.00 | 184.00 | 186.00 | 179.00 | 191.00 |
| Phonation | SNR | 2.30 | 0.83 | 1.29 | 1.87 | 4.91 | 3.45 | 64.09 | 11.62 | 0.31 | 0.37 | 3.27 | 4.89 |
|  | Jitter | 1.42 | 1.78 | 1.20 | 0.66 | 0.47 | 0.45 | 1.40 | 2.12 | 10.59 | 8.02 | 0.62 | 0.84 |
|  | Shimmer | 3.89 | 5.87 | 4.08 | 4.55 | 5.20 | 2.71 | 5.48 | 6.42 | 18.39 | 22.82 | 4.72 | 7.01 |
| Articulation | SNR | 0.27 | 0.28 | 0.49 | 0.65 | 0.33 | 0.30 | 0.84 | 0.51 | 0.79 | 1.21 | 0.59 | 0.75 |
|  | Jitter | 6.79 | 5.92 | 6.26 | 5.48 | 9.99 | 8.85 | 10.42 | 10.49 | 11.60 | 12.82 | 6.82 | 5.67 |
|  | Shimmer | 21.17 | 17.69 | 18.86 | 28.55 | 27.25 | 28.10 | 38.84 | 22.53 | 24.16 | 30.21 | 14.51 | 16.06 |
| Reading | SNR | 0.94 | 0.49 | 0.68 | 0.61 | 0.57 | 0.63 | 1.34 | 0.78 | 2.44 | 5.91 | 0.83 | 0.63 |
|  | Jitter | 7.66 | 6.34 | 5.82 | 7.99 | 8.15 | 7.20 | 7.50 | 6.38 | 16.27 | 13.41 | 7.23 | 5.76 |
|  | Shimmer | 15.42 | 12.91 | 15.31 | 13.90 | 15.88 | 15.00 | 15.75 | 16.72 | 18.67 | 19.10 | 15.47 | 15.58 |
| Balance | Watch X PL | 3.12 | 3.25 | 4.91 | 4.37 | 2.74 | 2.74 | 3.19 | 3.47 | 4.51 | 6.39 | 5.82 | 3.59 |
|  | Watch Y PL | 4.38 | 3.97 | 4.91 | 4.77 | 2.59 | 3.18 | 3.17 | 3.58 | 5.67 | 9.43 | 6.55 | 4.26 |
|  | Watch Z PL | 4.06 | 3.83 | 5.64 | 5.27 | 3.07 | 3.55 | 3.06 | 4.11 | 5.08 | 7.16 | 5.69 | 4.09 |
|  | Phone X PL | 2.63 | 2.79 | 4.92 | 3.15 | 2.67 | 1.89 | 2.33 | 2.08 | 3.03 | 3.23 | 3.47 | 3.08 |
|  | Phone Y PL | 1.96 | 1.99 | 3.11 | 2.98 | 1.98 | 1.73 | 2.04 | 1.92 | 2.92 | 2.76 | 2.66 | 2.45 |
|  | Phone Z PL | 1.80 | 2.05 | 4.82 | 2.87 | 2.45 | 1.90 | 2.07 | 1.96 | 3.68 | 3.78 | 3.26 | 2.58 |
| Gait | Watch X PL | 185.06 | 221.83 | 208.49 | 222.46 | 219.53 | 199.73 | 191.66 | 217.85 | 104.70 | 97.66 | 113.33 | 111.92 |
|  | Watch Y PL | 178.76 | 181.29 | 207.74 | 214.96 | 149.25 | 151.27 | 158.76 | 169.21 | 108.55 | 100.72 | 104.04 | 102.66 |
|  | Watch Z PL | 300.15 | 350.82 | 326.59 | 337.76 | 249.14 | 234.28 | 234.36 | 246.94 | 222.06 | 190.84 | 191.15 | 230.03 |
|  | Phone X PL | 243.36 | 289.51 | 305.27 | 396.54 | 276.09 | 307.55 | 284.90 | 341.43 | 273.91 | 191.50 | 204.74 | 248.23 |
|  | Phone Y PL | 376.41 | 410.86 | 563.04 | 581.12 | 330.13 | 290.44 | 313.47 | 363.16 | 335.26 | 262.80 | 251.70 | 319.18 |
|  | Phone Z PL | 456.12 | 463.48 | 466.51 | 493.92 | 341.37 | 351.91 | 316.66 | 362.17 | 335.38 | 262.41 | 268.13 | 324.89 |
| Arm Stability | Resting SD | 0.00 | 0.00 | 0.00 | 0.00 | 0.00 | 0.00 | 0.00 | 0.00 | 0.00 | 0.00 | 0.00 | 0.00 |
|  | Postural SD | 0.00 | 0.00 | 0.00 | 0.00 | 0.00 | 0.00 | 0.00 | 0.00 | 0.00 | 0.00 | 0.00 | 0.11 |
|  | Pronation SD | 0.03 | 0.03 | 0.05 | 0.05 | 0.01 | 0.01 | 0.02 | 0.05 | 0.14 | 0.19 | 0.03 | 0.12 |

- Fine motor skills trended towards reduced performance.
- Visual short-term memory (VSTM) trended towards longer response times.
- Gait and balance tasks trended towards altitude-related increase in accelerometer variability.
- Postural stability trended towards altitude-related increases in accelerometer variability. Fourier analysis of postural tremor task data demonstrated broadband increase in frequencies less than 10 Hz (Figure 3).
- All voice tasks (phonation, articulation, reading) demonstrated higher SNR at altitude.

## DISCUSSION

This equipment validation study was able to confirm the fidelity of microphone and accelerometer function of a smartphone at altitude. Limited prior data evaluating smartphone sensor performance at altitude is available, although the smartphone was successfully used to assess heart rate variability in cyclists at altitude [17]. To our knowledge this study is the first validation of high fidelity microphone and accelerometer data capture against known standards at sea level and altitude. Excellent correlation was noted between observed and expected frequencies under all conditions. This supports the use of smartphone sensors to assess a range of cognitive and physical functions in subjects exposed to high altitude.

Impairment of fine motor skills at altitude has been reported in military precision marksmen [18] and was reconfirmed in this study as significantly abnormal. Another military study described a reduction in cognitive performance at altitude [19]. Our study was consistent, showing that fine motor skills deteriorate at altitude. Conversely, we showed that visual short term memory is maintained at altitude but has longer response times. Neuropsychological assessment was used in airmen exposed to acute hypoxia in a simulated environment of 20,000 feet (6,100 m) of altitude to show that both memory encoding and response is abnormal, possibly reflecting more severe levels of hypoxia in that study [20]. Although hypoxia at similar altitudes using motor waveforms showed reduction in the motor functions of speech, we did not demonstrate a similar abnormality, possibly reflecting the different measurement systems used [21].

Hypoxia at lower levels of altitude is known to have effects on gait and balance in both normal subjects and those with chronic lung disease [2] [22] [23]. Similarly, postural control abnormalities have been reported in those trekking as high as 16,900 feet (5,150 m) [24]. Under the clinically significant condition of cerebral edema at altitude, a range of neurologic functions are impaired including gait [25]. We reconfirmed these observations, showing that both gait and balance are impaired with altitude-related increase in measurement variability. Our observation of postural tremor at altitude with a frequency range below 10Hz is consistent with tremor in the 6-12 Hz range previously reported in simulated acute hypoxia in healthy volunteers [26] [27]. The mechanism has been hypothesized as activated physiological tremor [26], but Parkinsonism has been reported at altitude [4], and Parkinson’s disease is consider a relative contraindication for travel to altitude [3]. The recent WATCH-PD study used the same software analysis (BrainBaseline ™) to successfully correlate symptoms in Parkinson’s Disease patients with a similar range of movement biomarkers at sea level [10]. Hypoxia may also be a key factor in the pathogenesis of Parkinson’s Disease [28].

This equipment validation study validated the use of microphone and accelerometer sensors (iPhone 14) at sea-level and 12,500 feet. We used a smartphone and smartwatch to reconfirm prior reports of the effects of acute hypoxia on cognition, speech, fine motor function, gait, and balance. Further studies could use expand on these biomarker observations by using smartphone and smart watch technology to assess high altitude physiology. Potential applications include the pathogenesis of altitude induced illness and deterioration of Parkinson’s Disease, as well as prediction of aircrew safety incidents and military performance.

